# *Saccharomyces* spores are born prepolarized to outgrow away from spore-spore connections and penetrate the ascus wall

**DOI:** 10.1101/2020.06.30.181362

**Authors:** Lydia R. Heasley, Emily Singer, Michael A. McMurray

## Abstract

How non-spore haploid *Saccharomyces* cells choose sites of budding and polarize towards pheromone signals in order to mate has been a subject of intense study. Unlike non-spore haploids, sibling spores produced via meiosis and sporulation by a diploid cell are physically interconnected and encased in a sac derived from the old cell wall of the diploid, called the ascus. Non-spore haploids bud adjacent to previous sites of budding, relying on stable cortical landmarks laid down during prior divisions, but since spore membranes are made de novo it was assumed that, as is known for fission yeast, *Saccharomyces* spores break symmetry and polarize at random locations. Here we show that this assumption is incorrect: *Saccharomyces cerevisiae* spores are born prepolarized to outgrow, prior to budding or mating, away from interspore bridges. Consequently, when spores bud within an intact ascus, their buds locally penetrate the ascus wall, and when they mate, the resulting zygotes adopt a unique morphology reflective of re-polarization towards pheromone, which we dub the derrière. Long-lived cortical foci containing the septin Cdc10 mark polarity sites, but the canonical bud site selection program is dispensable for spore polarity, thus the origin and molecular composition of these landmarks remain unknown. These findings demand further investigation of previously overlooked mechanisms of polarity establishment and local cell wall digestion, and highlight how a key step in the *Saccharomyces* life cycle has been historically neglected.

## 2. Introduction

Most steps in the *Saccharomyces cerevisiae* life cycle have been described in great detail at the cellular and molecular levels. For example, in the YeastBook series of reviews published by the journal *GENETICS*, forty three articles to date “span the breadth of *Saccharomyces* biology” (https://www.genetics.org/content/yeastbook). Here and in other sources can be found numerous mechanistic insights into the ways that budding yeast cells choose bud sites by positioning polarity factors according to mating-type-specific cortical landmarks; how a potent extrinsic signal, mating pheromone, can override these landmarks; and how cells use hydrolytic enzymes to remodel their cell walls upon budding and cell fusion. The unique morphology yeast cells adopt as they grow chemotropically toward a pheromone source even garnered an enduring and endearing name, the shmoo, based on its resemblance to a comic strip character from the mid 1900s. Indeed, early studies of the signal transduction cascades underlying the pheromone response and the ways that yeast cells repolarize to track pheromone gradients were foundational in our general understanding of eukaryotic signal transduction and polarity determination.

Accordingly, we find it remarkable how little is known about a step of the *Saccharomyces* life cycle for which polarity, chemotropism, and cell wall remodeling are crucial. Germination is the process by which spores awaken from a nearly dormant state, break out of a rigid, specialized cell wall, and grow in a single direction by producing new membrane and cell wall. Only a single review dedicated to germination has been published, over a decade ago (Geijer et al., 2010).

Since most natural isolates of *S. cerevisiae* are diploid, most spores are haploid, and germinating spores are capable of mating immediately (without a prior budding event) if a suitable mating partner is sufficiently close by (Joseph-Strauss et al., 2007). The assumption from the literature is that mating between spores follows the same rules as what has been worked out for non-spore haploids, but this has not been rigorously tested.

Alternatively, and often regardless of the proximity of a mating partner (McClure et al., 2018), following outgrowth haploid spores can enter S phase and bud from the tip of the outgrowth. The most recent comprehensive review of cell polarity in yeast states that “in yeasts, germinating spores lack obvious positional cues and appear to break symmetry to initiate polar growth” (Chiou et al., 2017). This statement is clearly true for the fission yeast *Schizosaccharomyces pombe*, where an elegant study demonstrated that polarity factors, including active Cdc42, wander randomly around the cortex of growing spores until the outer spore wall breaks, whereupon polarity is stabilized at the site of wall rupture (Bonazzi et al., 2014). In *S. cerevisiae*, on the other hand, a single polarity site is already apparent in dormant spores, prior to germination; the only known marker of this site is the septin protein Cdc10 (Joseph-Strauss et al., 2007). Polarization of the actin cytoskeleton, a prerequisite for outgrowth, occurs only after germination begins (Kono et al., 2005). In non-spore cells, septins are recruited to the site of future budding by active Cdc42 and Cdc42-interacting proteins (Iwase et al., 2006); it is not known how Cdc10 is deposited during sporulation at a single site on each spore. During sporulation the spore cortex is synthesized de novo, precluding any influence on bud site selection by persistent cortical landmarks produced by previous budding events.

Yeast sporulation takes place within a single (usually diploid) cell and generates (usually four) spores encased within the original cell wall of the sporulating cell, called the ascus (Neiman, 2011). Whereas in *S. pombe* the ascus wall is globally digested by cell wall hydrolases immediately upon the successful completion of meiosis (Dekker et al., 2007; Encinar del Dedo et al., 2009; H. Guo & King, 2013), in *Saccharomyces* the ascus wall remains intact throughout sporulation. In the absence of external factors, such as those found in insect guts (A. E. Coluccio et al., 2008), the *Saccharomyces* ascus wall breaks down slowly during germination, often persisting long enough for spores to mate and bud following germination. *Saccharomyces* spores (but not spores of *S. pombe*) are also interconnected by cell wall junctions called interspore bridges that persist following ascus wall digestion (A. Coluccio & Neiman, 2004). Outgrowth by germinating spores first requires local break down of a rigid outer spore wall that confers stress resistance via layers of chitosan and polymerized dityrosine (in *Saccharomyces* (Briza et al., 1990)) or a proteinaceous coat (in *S. pombe* (Fukunishi et al., 2014)). Subsequent budding or mating requires local breakdown of the newly-emerged cell wall at the site of budding or fusion, respectively. If *Saccharomyces* spores bud when the ascus wall is still intact, those buds can locally penetrate the ascus wall, as visualized by electron microscopy many decades ago (Hashimoto et al., 1958; Rij, 1978; Sando et al., 1980). One of us (M.M.) noticed this phenomenon using light microscopy in the context of a more recent study, and verified that the protrusions were buds by visualizing bud necks marked with fluorescently tagged septin proteins (McMurray & Thorner, 2008). The mechanisms by which germinating *Saccharomyces* spores digest cell walls during outgrowth, budding and mating have not been investigated.

Here we describe how the sites where spores bud and fuse upon mating, and the localization of Cdc10 with relation to the position of other haploid spores produced by the same diploid mother, demonstrate that each *S. cerevisiae* spore is born pre-polarized to direct outgrowth away from its “sibling” spores. Spore buds are thereby positioned to penetrate the ascus wall.

## 3. Materials and Methods

### 3.1. Yeast strains and media

#### Stable cortical foci of the septin Cdc10 at the ascus periphery

All strains used in the new experiments described here are of the S288C strain background, specifically derived from the “designer deletion” strains BY4741 (*MAT***a***his3Δ1 leu2Δ0 met15Δ0 ura3Δ0*), BY4742 (*MATα his3Δ1 leu2Δ0 lys2Δ0 ura3Δ0*), and BY4743 (the diploid formed by mating BY4741 and BY4742). Unless specified otherwise in the figure legend, all experiments were done with a diploid strain made by mating FY2742 (*MATα his3Δ1 leu2Δ0 lys2Δ0 ura3Δ0 TAO1 MKT1 RME1*) and FY2839 (*MAT***a** *his3Δ1 leu2Δ0 lys2Δ0 ura3Δ0 TAO1 MKT1 RME1*), two haploid strains that carry at three loci (*TAO1*, *MKT1*, and *RME1*) dominant alleles from the efficiently-sporulating SK-1 strain background (Kloimwieder & Winston, 2011). JTY3985 carries *CDC10-GFP* integrated at the *CDC10* locus using the *URA3* selectable marker and is otherwise isogenic to BY4741 (Johnson et al., 2015). MMY0341 (*gas1Δ::kanMX /gas1Δ::kanMX*) MMY0286 (*acf2Δ::kanMX, acf2Δ::kanMX*), MMY0291 (*rsr1Δ::kanMX, rsr1Δ::kanMX*) were retrieved from the homozygous diploid deletion collection derived from BY4743 (Giaever et al., 2002). Previously published Gas1-GFP localization data reproduced here used a diploid strain of the SK-1 background in which both genomic copies of *GAS1* were deleted and Gas1-GFP was expressed from a high-copy plasmid (Rolli et al., 2011).

Haploid strains were mated together using sterile toothpicks by mixing approximately equal amounts on the surface of YPD agar (per liter: 10 g yeast extract, 20 g peptone, 20 g dextrose, 2% agar) in a petri dish. After overnight incubation at 30°C, cells from the mixture were streaked with a toothpick to agar media that was either selective for the diploid, or non-selective (YPD). In the latter case, diploid clones were identified as individual colonies containing cells that were able to sporulate.

To induce sporulation, diploid cells were cultured in 5 mL liquid YPD (per liter: 10 g yeast extract, 20 g peptone, 20 g dextrose) in glass culture tubes rotated in a roller drum overnight to near-saturation. For analysis of *rsr1Δ/rsr1Δ* cells in “old” rich medium, the culture time in YPD was extended for 6 additional days. A 200-μL aliquot of these cells was washed with 5 mL sterile water and resuspended in 2.5 mL sporulation medium (1% potassium acetate, 0.05% glucose, 20 mg/L leucine, 40 mg/L uracil) in a new tube to an optical density at 600 nm of approximately 0.5. These cell suspensions were rotated at 22°C for at least 4 days, after which time the percentage of cells that had formed mature asci did not noticeably increase. To induce germination, aliquots of cells from sporulation cultures were pelleted and resuspended in 1 mL YPD in 1.7-mL microcentrifuge tubes and rotated at 22°C or 30°C for 4-6 hours (or as indicated in figure legends). To digest the ascus wall, a 50-μL aliquot of cells from sporulation culture was pelleted and resuspended in 1 mg/mL Zymolyase-20T (#320921, MP Biomedicals) dissolved in water and incubated at 30°C for 10 minutes.

### 3.2. Fluorescence labeling

FM™ 4-64 dye (N-(3-Triethylammoniumpropyl)-4-(6-(4-(Diethylamino) Phenyl) Hexatrienyl) Pyridinium Dibromide, from Molecular Probes, Inc. # T3166) was dissolved in dimethyl sulfoxide to make a 1.6 μM stock solution. An aliquot of cells from a sporulation culture that had been in sporulation medium for 18 hours was pelleted and resuspended in 50 μL ice-cold YPD to which 1 μL of the dye stock was then added. After 20 minutes on ice in the dark, the cells were pelleted again and washed twice by resuspension in 1 mL ice-cold water each time. Finally, the cells were suspended in 50 μL water and visualized by microscopy using the Texas red LED filter cube.

Calcofluor white M2R was dissolved in water at a concentration of 10 mg/mL. 1 μL was added to a 100-μL aliquot of cells that had been in sporulation medium for 4 days, after which the cells were washed three times with water by pelleting and resuspension. After the third wash, the cells were resuspended in 1 mL YPD and incubated at 30°C for 6 hours, then pelleted and resuspended in 50 μL before imaging.

### 3.3. Microscopy and imaging

Aliquots of cells from sporulation cultures were imaged directly on agarose pads made with 1% agarose in water or, in the case of cells expressing Cdc10-GFP, before and after a 3.5-hour interval of incubation at 30°C on an agarose pad made with 1% agarose in sporulation medium. Germinating cells were pelleted and resuspended in water before applying to agarose pads. All images were captured on an EVOSfl all-in-one microscope (ThermoFisher Scientific, Waltham, MA) with an Olympus 60× Plan-Apo oil objective (numeric aperture 1.42). Filter cubes were as follows: GFP (AMEP4651, excitation 470/22 nm, emission 510/42 nm), Texas red (AMEP4655, excitation 585 nm, emission 624 nm), RFP (AMEP4652, excitation 531/40 nm, emission 593/40 nm), and DAPI (AMEP4650, excitation 357/44 nm, emission 447/60 nm). Images were cropped and adjusted (always the same way for each image of the same type from the same experiment), and inverted in Photoshop (Adobe, San Jose, CA).

## 4. Results

### 4.1. Stable cortical foci of the septin Cdc10 at the ascus periphery

That single *S. cerevisiae* spores are born pre-polarized was known from the discrete localization of the septin Cdc10 to a single site on the spore cortex from which cell wall outgrowth occurs (Joseph-Strauss et al., 2007). However, it was not known how the site of spore pre-polarization relates to the spatial relationship between the spores, the interspore bridges, and the ascus wall. If spore buds commonly penetrate the ascus wall, then spores should be polarized to outgrow and bud away from the interspore bridges at the center of the ascus. To test this prediction, we visualized Cdc10-GFP in spores within intact, mature asci. As is true of diverse fluorescently-tagged proteins, strong signal was observed in the areas between the spores (Figure 1A), presumably reflecting Cdc10-GFP molecules that were outside the prospore membranes as they closed and thus remained in the ascal cytoplasm, which becomes concentrated between the spores as the ascus wall compresses tightly to surround the spores during the final stages of ascus maturation. Cdc10-GFP signal is not found at these locations in spores that are isolated from asci (Joseph-Strauss et al., 2007). Apart from this signal, single Cdc10-GFP puncta were found in many (but not all) spores (Figure 1A); the failure to observe a punctum in every spore may reflect the fact that only one allele of *CDC10* in the diploid strain expresses the GFP fusion. While it is known that all spores from such *CDC10-GFP/CDC10* heterozygous diploids inherit some pre-existing Cdc10-GFP protein, regardless of their haploid genotype, those that fail to inherit the *CDC10-GFP* allele have fainter signal (Joseph-Strauss et al., 2007). When they were visible, Cdc10-GFP puncta were found in locations consistent with a mode of outgrowth upon germination that precedes spore budding through an intact ascus wall (Figure 1A and data not shown).

**Figure 1.**
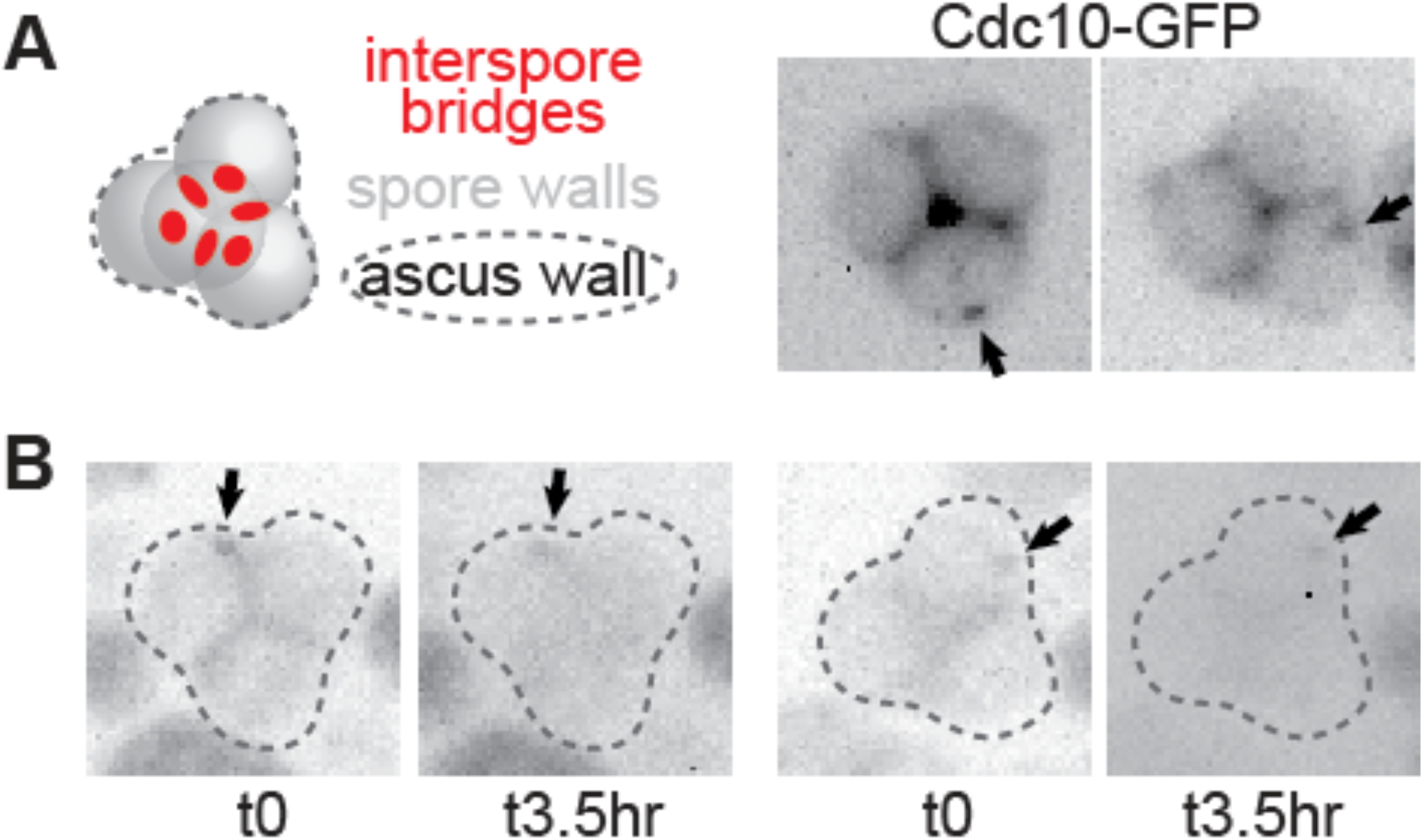
Stable cortical foci of the septin Cdc10 persist on the periphery of asci. (A) At left is an illustration of a typical pyramidal four-spored ascus in which the three spores at the “base” of the pyramid are in the same focal plane. Red circles indicate interspore bridges. At right, Cdc10-GFP fluorescence in intact asci from sporulation culture. Arrows indicate cortical foci. (B) As in (A), but the same asci were visualized before and after a 3.5-hr interval on solid sporulation medium. Cells are of diploid strain made by mating haploid strains JTY3985 and FY2742.

Spores can survive for long periods in nutrient-free conditions and then, once nutrients are provided, germinate efficiently. We see no obvious effect on the site of budding during germination of the length of time between sporulation and germination (unpublished observation); indeed, the images in Figure 1A were taken of asci that had been in sporulation medium for over 7 days, with the vast majority of asci in these cultures having visibly completed maturation by day 4. If Cdc10 foci mark the location of spore pre-polarization, then these cortical foci must not diffuse to any great extent over time. Alternatively, they may be able to diffuse on regions of the cortex far from the interspore bridges, but may be prevented from diffusing into the cortical areas near interspore bridges. To distinguish between these possibilities, we visualized Cdc10-GFP foci before and after an interval of 3.5 hrs. We could detect no change in location during this time (Figure 1B). These data suggest that diffusion of the cortical Cdc10-GFP foci is highly restricted, potentially allowing spores to maintain cortical polarity for long periods of time.

### 4.2. Ascus wall penetration upon spore budding points to prepolarization of spores away from interspore bridges

If spores are pre-polarized to outgrow upon germination towards the periphery of the ascus, then when germinating spores bud while they are still within the ascus, they should locally penetrate the ascus wall, such that the buds protrude from the ascus surface. If, instead, spore pre-polarization is random, then spore budding should frequently result in budding in between the spores, within the ascus, without penetrating the ascus wall. We exploited the weak red autofluorescence of the post-germination spore wall (Joseph-Strauss et al., 2007) to identify trapped buds (non-fluorescent cell wall outgrowth) within the crowded environment of an intact ascus full of germinating spores. Among hundreds of such asci we have only seen a single likely case of a bud “trapped” within the ascus (Figure 2A).

**Figure 2.**
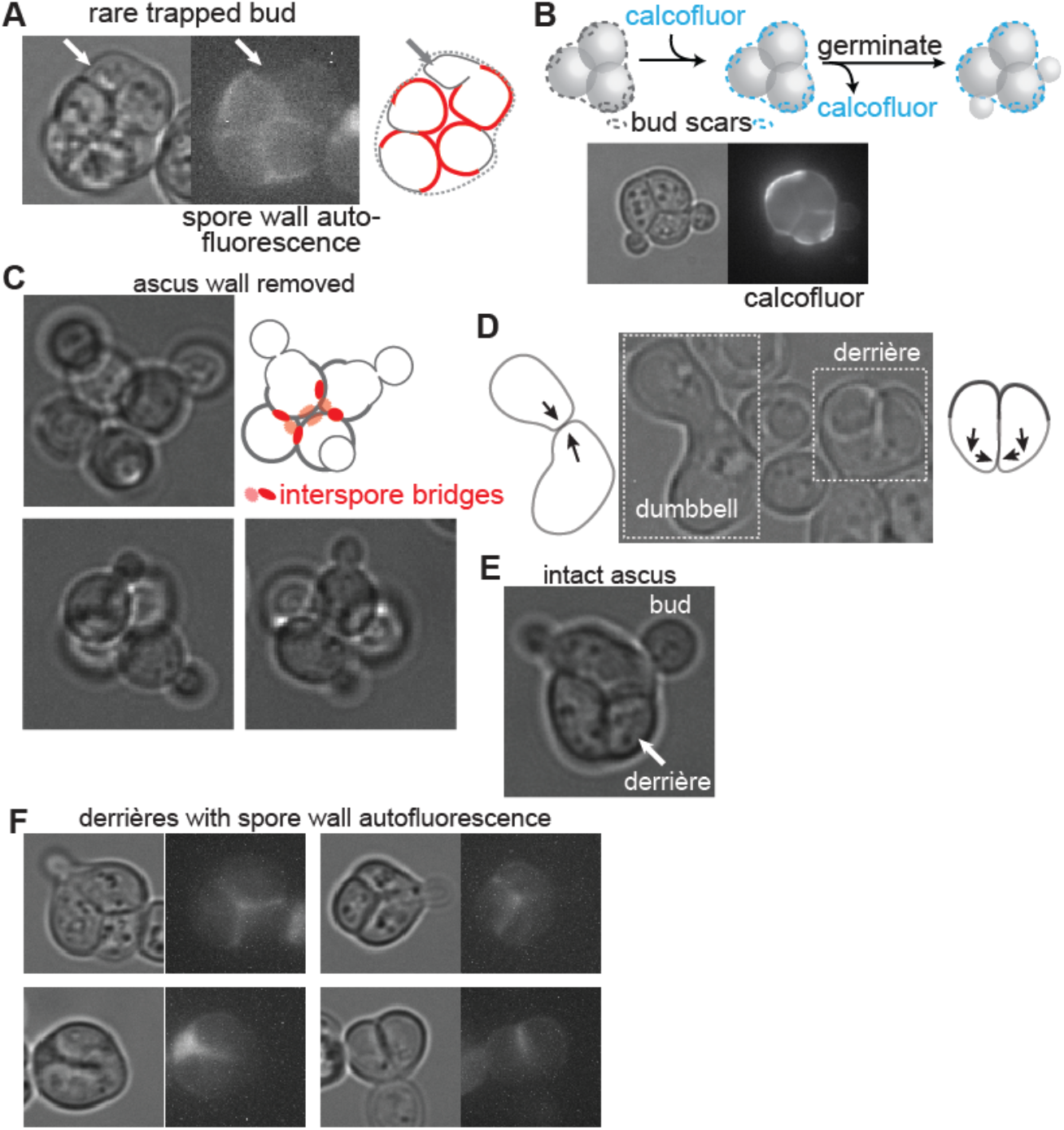
*Saccharomyces* spores outgrow away from interspore bridges upon germination. (A) Germinating ascus as viewed by transmitted light and with autofluorescence of the spore wall visualized with an RFP filter. (B) According to the illustration, asci were exposed to the chitin-binding dye calcofluor white and then, after washing away free dye, allowed to germinate. Pre-existing bud scars on the ascus wall were deposited during diploid budding events prior to sporulation. Calcofluor fluorescence and transmitted light are shown. (C) Tetrads for which the ascus wall was removed by exposure to Zymolyase prior to germination. In the illustration red circles are interspore bridges, and the spore wall is thicker than the new, vegetative cell wall. (D) Image taken several hours after asci were allowed to germinate, showing dumbbell-shaped zygote and derrière-shaped zygote, with illustrations of presumptive directions of outgrowth prior to fusion. (E) Germinating ascus showing two buds penetrating the ascus wall and the other two spores fused into a derrière. (F) As in (A), after 7.75 hours of germination, showing localization of the autofluorescent spore wall with regard to the shape of the derrière.

As an independent way to visualize bud penetration of the ascus wall, we performed a pulse-chase with calcofluor white, which fluorescently labels cell wall chitin (Cabib & Bowers, 1975). The ascus wall in mature asci was labeled with calcofluor, and then excess dye was washed away prior to the induction of germination by addition of rich (YPD) medium (Figure 2B). New cell wall synthesized in the absence of calcofluor white is non-fluorescent. Spore buds emerged from holes in the ascus wall, which otherwise remained intact (Figure 2B). Thus budding by germinating spores is directed away from the interspore bridges that connect spores to each other, and towards the ascus wall that surrounds them. Budding away from interspore bridges was also apparent for interconnected spores in which the ascus wall was enzymatically removed (Figure 2C).

### 4.3. Unique zygote morphology provides independent evidence of spore prepolarization

We noticed independent evidence of spore pre-polarization in the morphology of zygotes produced by spores that mated within the ascus. Figure 2D shows an image of cells from a population of germinating asci in which a zygote formed by non-spore haploid cells (or possibly a spore with a non-spore cell) is adjacent to a zygote formed by mating between sibling spores. The dumbbell morphology of the zygote formed by non-spore mating is consistent with the morphology established in the literature for mating between wild-type haploids (Sena et al., 1973), where prior to cell fusion each partner grows directly towards the other, following a pheromone gradient. The zygote formed from intra-ascus mating between spores is distinctly different, as if both mating partners initially grew in approximately the same direction, roughly perpendicular to a line directly connecting them, and then redirected growth towards each other prior to fusion (Figure 2D). Similar to the use of the term “shmoo” to refer to the unique morphology adopted by non-spore cells just prior to mating, we sought a new term to refer to the unique morphology of zygotes produced by mating between sibling spores. Inspired by the inescapable resemblance of one side of the resulting shape to the shape of the human buttocks, we propose the term “derrière”. Derrières were also seen within intact asci after germination (Figure 2E).

The rigid outer spore wall is likely a barrier to outgrowth, which requires cell wall breakdown at the site of fusion. Cell wall expansion upon germination is thus restricted to the site of outer spore wall breakdown, and the residual spore wall changes little upon germination and thereafter. If spores break down the outer spore wall at sites approximately opposite from the interspore bridges, then two adjacent spores within an ascus should grow along vectors that do not converge and instead are parallel or, more likely, divergent (Figure 2D). If redirected growth towards a pheromone source is only possible for vegetative cell wall, where polarized exocytosis targets cell wall synthesis enzymes to specific sites, then some amount of cell wall outgrowth is presumably required before that cell wall outgrowth can be redirected towards the pheromone source. This model predicts derriére-shaped zygotes with spore wall in the twin bulges. To test this model, we visualized spore wall autofluorescence within derrières. As expected, spore wall fluorescence was restricted to the twin bulges of derrières (Figure 2F). Buds then emerged from the site of fusion (Figure 2D,F).

### 4.4. The old plasma membrane of the sporulating diploid cell disappears prior to germination

In addition to local digestion via budding, the ascus wall breaks down globally during germination, albeit on a slower time scale. In *S. pombe*, two hydrolytic enzymes, the α-1,3-glucanase Agn2 and the endo-β-1,3-glucanase Eng2, reside in the ascal cytoplasm because they lack signal sequences to drive secretion (Dekker et al., 2007; Encinar del Dedo et al., 2009). It has been speculated that after completion of spore wall synthesis, the old plasma membrane of the diploid fission yeast cell “may disintegrate through an unknown mechanism” (Dekker et al., 2007) allowing the two enzymes access to their substrates.

If in *Saccharomyces* spore buds frequently penetrate the ascus wall, we wondered if in doing so they also penetrate the old plasma membrane of the sporulating cell, or if, as is thought for fission yeast, that membrane has already been destroyed. Late in sporulation, vacuole lysis releases hydrolytic enzymes that destroy, among other things, nuclei that were not protected by prospore membrane engulfment (Eastwood et al., 2012). The functional integrity of old plasma membrane becomes compromised at the same time, suggesting that this membrane may also be destroyed by vacuole lysis (Eastwood & Meneghini, 2015). A direct examination of plasma membrane persistence during sporulation has not, to our knowledge, been reported. To label the plasma membrane in cells undergoing sporulation, we exposed an asynchronously sporulating culture to FM4-64, a lipophilic dye (Vida & Emr, 1995). As can be seen in Figure 3, very early in sporulation, when spore walls had not yet been made, only the plasma membrane was labeled. Slightly later, when spore walls had been made but the ascus wall had not yet compressed around the spores, the old plasma membrane was labeled and, in some cases, the spore membranes were also labeled. In mature asci, in which the ascus wall had compressed tightly around the spores, only spore membrane labeling was visible. We interpret these results as an intact plasma membrane “shielding” the spore membranes from labeling early in sporulation, then losing structural integrity later in sporulation, and finally disappearing altogether in mature asci. If our interpretation is correct, when spores germinate and bud within an intact ascus, they only penetrate an old cell wall, and not an old membrane, in order to exit the ascus.

**Figure 1.**
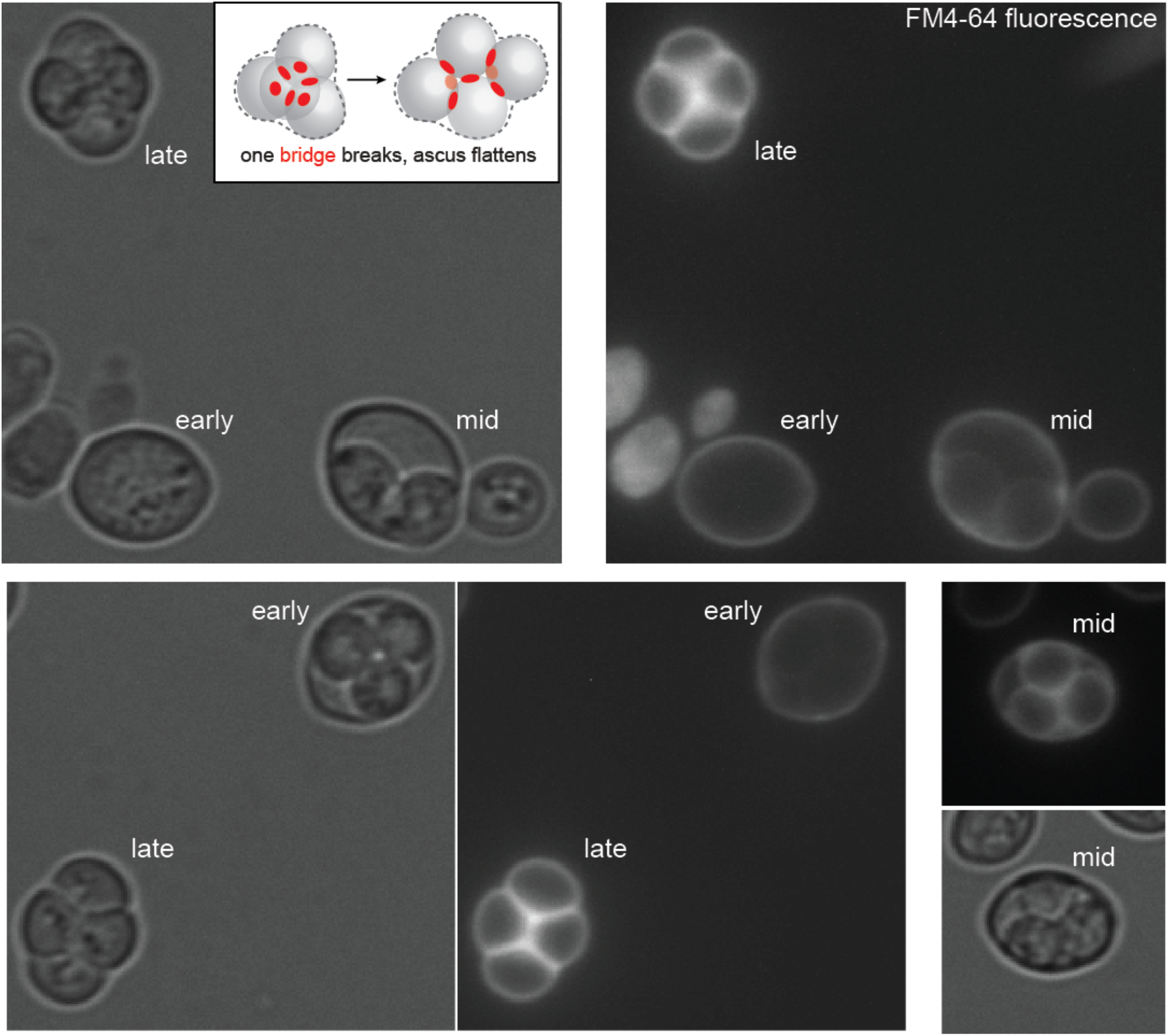
Gradual loss of the old plasma membrane during sporulation. An asynchronously sporulating culture was exposed briefly to the lipophilic dye FM™4-64. The stage of sporulation is labeled and was estimated based on spore and ascus wall appearance by transmitted light. Inset, an illustration of the arrangement of spores in a tetrahedral ascus with all six interspore bridges (red circles) intact, and in a rhomboid ascus in which one of the bridges has broken (dashed red circle) as an ascus flattens under a coverslip.

### 4.6. Global ascus wall digestion does not require the putative cell wall hydrolase Acf2

In the *S. cerevisiae* genome, the *ACF2* gene (alias *ENG2*) encodes a homolog of *S. pombe* Eng2, one of the enzymes responsible for global ascus wall digestion (Encinar del Dedo et al., 2009). Like Eng2, Acf2 lacks a predicted signal sequence. In *S. pombe*, the absence of either Agn2 or Eng2 is sufficient to almost completely prevent global ascus wall digestion, resulting in mature spores trapped within an ascus wall (Dekker et al., 2007; Encinar del Dedo et al., 2009). If in *S. cerevisiae* Acf2 digests the ascus wall during germination, we predicted that it should either be translated near the end of sporulation and then secreted into the ascal cytoplasm upon germination, or both translated and secreted immediately upon germination. No translation data are available for germination, but existing ribosome profiling data from synchronously sporulating cells (Brar et al., 2012) demonstrate that Acf2 translation spikes at the last stage of sporulation, just before mature spores are produced (Figure 4A). If gradual Acf2-mediated digestion gradually thins the ascus wall during germination, we predicted that in asci lacking Acf2, the ascus wall might remain too thick and inhibit penetration by spore buds. However, we saw no discernible difference in the frequency or morphology of instances in which the buds of *acf2Δ* spores penetrated the ascus wall upon germination (Figure 4B). In *S. cerevisiae*, either Acf2 performs a different function than Egn2, or loss of a single digestive enzyme is not enough to toughen the ascus wall to an extent that is impenetrable by buds.

**Figure 2.**
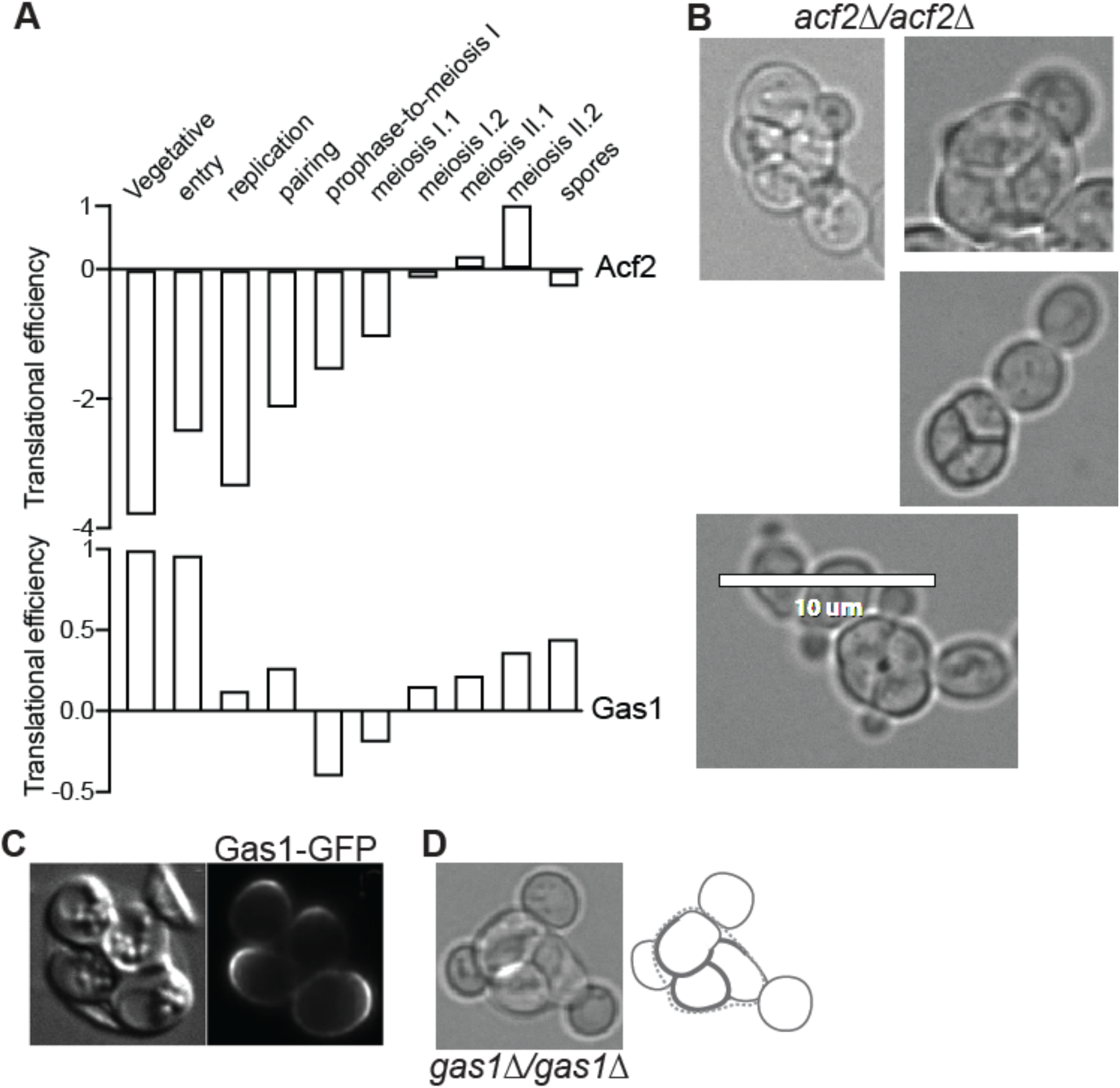
Penetration of the ascus wall by spore budding does not require Gas1 or Acf2. (A) Translational efficiencies (a measure of ribosome occupancy per mRNA) for Acf2 and Gas1 at various timepoints during sporulation. Data are from (Brar et al., 2012). (B) Germinating asci of *acf2Δ/acf2Δ* strain MMY0286 showing penetration of the ascus wall by spore budding. Scale bar, 10 μm. (C) Localization of Gas1-GFP to sites of spore outgrowth upon germination. Reproduced from (Rolli et al., 2011). (D) A germinating ascus of *gas1Δ/gas1Δ* strain MMY0341.

### 4.5. Local ascus wall digestion does not require the putative cell wall hydrolase Gas1

To identify candidate enzymes that may mediate local digestion of the ascus wall during germination, we searched the literature for published localization patterns of cell wall hydrolytic enzymes, with the logic that a protein responsible for local wall digestion should localize at the site of digestion. Translation of the β-1,3-glucanosyltransferase Gas1 translation increases at the end of sporulation (Figure 4A), and while this pattern was not specifically noted by the authors, examination of a GFP-tagged allele of Gas1 in spores germinating within an intact ascus revealed that Gas1-GFP clearly localizes to a broad region of the spore cortex opposite from interspore bridges (Rolli et al., 2011) (Figure 4C). To ask if Gas1 is required for the ability of buds to penetrate the ascus wall, we monitored germination by *gas1Δ/gas1Δ* mutant diploid cells. Consistent with the phenotype reported in the literature for non-spore haploid *gas1Δ* cells (Watanabe et al., 2009), the buds produced by *gas1Δ* spores were oddly-shaped, but they had no problems penetrating the ascus wall (Figure 4D). Thus deletion of *GAS1* is insufficient to prevent local ascus wall digestion by germinating spores.

### 4.6. The canonical bud site selection pathway does not drive polarity in spores

Spores polarize approximately opposite from the cluster of interspore bridges that connect them to the other spores within in ascus. Similarly, diploid non-spore cells usually polarize opposite from the previous bud site (Chiou et al., 2017). We noticed that two genes required for bipolar bud site selection by non-spore cells, *RSR1* (Bender & Pringle, 1989) and *SPH1* (Roemer et al., 1998), are transcriptionally induced early in germination (Joseph-Strauss et al., 2007). We wondered if the same pathway that controls bipolar budding in non-spore diploids also controls spore polarization. We sporulated *rsr1Δ/rsr1Δ* diploids and monitored budding upon germination.

Analysis of budding was complicated by the fact that many of the mutant cells had multiple buds even prior to germination (Figure 5A). Unlike mitotic DNA replication, premeiotic DNA replication is usually uncoupled from bud emergence. The few singly-budded asci found in nominally wild-type strain backgrounds presumably arise from diploid cells that were in a small window of early S phase when sporulation began and proceeded directly into meiosis upon completion of DNA replication (Croes et al., 1976). In the *rsr1Δ/rsr1Δ* mutants, however, either buds formed during multiple prior cell cycles failed to separate from the mother, or multiple buds formed simultaneously upon premeiotic S phase entry. We favor the latter interpretation, considering that simultaneous multiple budding is a known phenotype of *rsr1Δ* haploid cells expressing a synthetic fusion of the polarity scaffold protein Bem1 to the v-SNARE Snc2, and that this phenotype is exacerbated when cells are cultured in minimal, as opposed to rich, medium (Howell et al., 2009). The starvation conditions induced by sporulation medium may bypass the need for the Bem1-Snc2 fusion in driving multiple budding events. Indeed, we found that starving *rsr1Δ/rsr1Δ* diploid cells by prolonged culture (1 week) in rich medium, which does not induce sporulation, was sufficient to induce a bi-budded phenotype (Figure 5B).

**Figure 3.**
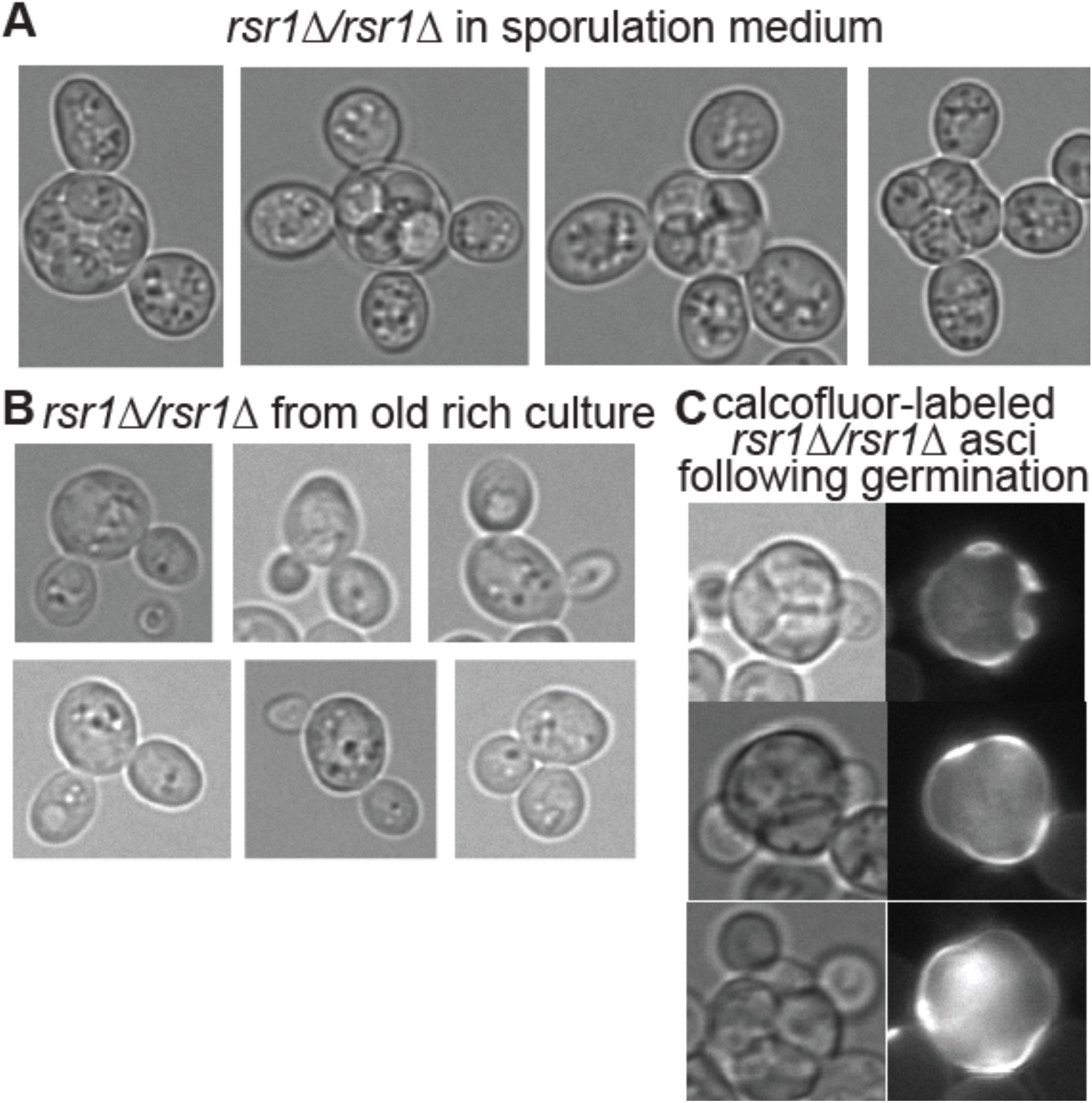
Cells lacking the polarity factor Rsr1 form multiple buds upon starvation but spore buds are able to penetrate the ascus wall. *rsr1Δ/rsr1Δ* strain MMY0291 was cultured in (A) sporulation medium, (B) rich medium for 1 week, or (C) sporulation medium, followed by pulse labeling with calcofluor white, and then rich medium to induce germination. Fluorescence images show calcofluor fluorescence.

To unambiguously identify buds produced during germination, we applied the calcofluor pulse-chase method. The vast majority of *rsr1Δ/rsr1Δ* asci that were not budded prior to germination showed buds penetrating the ascus upon germination (Figure 5C and data not shown). We conclude from these results that spore polarity is not determined by the canonical pathway that drives polarity during budding by non-spore cells.

## 5. Discussion

The evidence for prepolarization of *S. cerevisiae* spores has been hiding in plain sight for decades. For every published image we could find in which the orientation of a germinating spore relative to its sibling spores is discernible, the direction of outgrowth, budding and/or mating is consistent with prepolarization away from the interspore bridges. In *S. pombe* spores, on the other hand, lack of prepolarization has been clearly demonstrated (Bonazzi et al., 2014), and an assumption that *S. cerevisiae* is the same is understandable, given the many similarities with regard to mechanisms of sporulation. Several key differences are worth noting, however. Fission yeast spores lack interspore bridges, and ascus wall breakdown is a programmed event taking place immediately following successful completion of meiosis (H. Guo & King, 2013). Hence *S. pombe* spores are designed for dispersal, consistent with a mostly haploid lifestyle in this species, whereas *Saccharomyces* species appear to have evolved to prioritize return to diploidy following meiosis. From this perspective, why not prepolarize a spore directly towards its meiotic siblings, at least two-thirds of which will be of compatible mating type? A somewhat trivial explanation may be that interspore bridges represent a physical barrier to outgrowth, analogous to the difficulties that non-spore yeast cells encounter when mutations drive them to re-bud through a chitin-rich bud scar (Tong et al., 2007). We propose another model. In a crowded four-spored ascus, navigating gradients of multiple pheromones in order to identify a compatible partner is non-trivial (Rappaport & Barkai, 2012). If germinating spores grew into the center of the ascus, the situation would become even more complex, especially if spores budded and then switched mating types. Hence, prepolarization away from the center may facilitate pheromone sensing by simplifying the gradients of signals spores must navigate to find a mate.

For other septin functions, e.g., cytokinesis, Cdc10 acts together with other septin proteins as a stable hetero-oligomeric protein complex that act as diffusion barriers and scaffolds for the recruitment and cortical retention of other proteins (Oh & Bi, 2011). Polymerization of hetero-oligomeric septin complexes into filaments is required for cytokinesis (McMurray et al., 2011); no single septin is sufficient. Hence while we have not yet asked if other septins co-localize with Cdc10 to cortical foci in spores, we speculate that they do. What specific function, if any, Cdc10 might perform at these foci is an intriguing question for future inquiry. In non-spore cells, septins encircle active Cdc42 at both the site of bud formation (Okada et al., 2013) and the shmoo (Kelley et al., 2015), but the cortical puncta in spores are not obviously ring-shaped, and it is not known if Cdc42 (and/or another Rho-family GTPase) is also there.

How does Cdc10 arrive at this location? Cdc10 and other septins localize around the spindle pole bodies at the end of meiosis metaphase II, where the prospore membrane first appears, and then localize as bars and horseshoes associated with the prospore membrane as it grows (Pablo-Hernando et al., 2008). The nuclear envelope and prospore membrane are connected via the spindle pole body until the meiosis-specific components of the “meiotic outer plaque”, Mpc54 and Mpc70, are destroyed just after meiosis II (Knop & Strasser, 2000). The exocyst complex, which targets exocytic vesicle docking and fusion, is found (along with septins) at the meiotic outer plaque (Mathieson et al., 2010), at the shmoo tip (Kelley et al., 2015), and at the site of future budding by non-spore cells (W. Guo et al., 2001). Meiotic anaphase II pushes the spindle pole bodies toward the periphery of the ascus. If septins and/or the exocyst persist on the spore membrane at the former site of contact with the meiotic outer plaque, these “landmarks” will be near the periphery of the ascus, with the sites of prospore membrane fusion (and interspore bridges) in the center. Provided cortical diffusion of the landmarks is limited in mature spores, these sites would correspond to the Cdc10 localization patterns and sites of outgrowth that we observe upon germination. Future work will be required to test this model.

Ascus wall penetration by buds has been documented previously (Hashimoto et al., 1958; Rij, 1978; Sando et al., 1980) but the implications for spore polarity were not considered. The mechanism of local ascus digestion upon spore budding also remains unknown. Budding itself requires cell wall digestion at a single site on an unbudded cell. Thus when a spore buds inside an ascus, two walls are locally digested: the wall of the spore and the ascus wall. Another step in the yeast life cycle also requires local digestion of two walls: cell fusion upon mating. Here it is thought that targeted secretion of enzymes followed by cell-contact-limited diffusion restricts digestion to a narrow pore (Huberman & Murray, 2014). We imagine that a similar mechanism, perhaps involving the same enzymes, mediates ascus wall penetration during budding. Our results show that, despite localizing to the right place at the right time, Gas1 is dispensable for this process; other enzymes may act redundantly.

Intra-ascus mating events do not locally digest the ascus wall because repolarization towards the mating partner directs the digestive enzymes elsewhere. On the other hand, during inter-ascus mating events *Saccharomyces* spores penetrate four walls in order to fuse. Crowding promotes inter-ascus mating (Murphy & Zeyl, 2010), which emphasizes the importance of considering where germination occurs in nature and thus how evolution acted upon it. Here we know very little. Sporulation is most frequent at the colony periphery (Purnapatre & Honigberg, 2002), but to what extent wild yeast grow in such colonies is unknown.

Indeed, while studies of polarity determination and mating by isolated non-spore *Saccharomyces* cells in the laboratory setting have provided numerous insights into basic biology, considering what we now know about the *Saccharomyces* life cycle outside the lab (Tsai et al., 2008), budding and mating by non-spore haploids may represent a kind of backup plan for circumstances in which a spore is physically separated from, or chooses to ignore (McClure et al., 2018), its meiotic siblings. Even the spore-based proposed rationale for axial budding by haploids – to place two cells of one mating type adjacent to two cells of opposite mating type following mating-type switching by an isolated spore (Gimeno & Fink, 1992) – can now be viewed in an additional light: axial budding would help ensure that if a spore is still within an intact ascus when it buds a second time, the second bud will also penetrate the ascus wall. Our work thus highlights outstanding questions and lays a foundation for future studies.

## Acknowledgements

This work was supported by the National Science Foundation (award 1928900 to M.M.) and by the National Institute of General Medical Sciences of the National Institutes of Health (award T32GM008730 to L.R.H., as part of the Molecular Biology PhD Program). The content is solely the responsibility of the authors and does not necessarily represent the official views of the National Science Foundation or the National Institutes of Health.

## Notes

### Competing Interest Statement

The authors have declared no competing interest.

